# Osteoconductive material of a newly developed resin modified glass ionomer cement containing bioactive glasses for biomedical purposes

**DOI:** 10.1101/2020.05.27.118521

**Authors:** Ika Dewi Ana

## Abstract

Hybridization of resin modified- glass ionomer cement (RMGIC) and bioactive glass (BAG) may result higher mechanical strength and resistance to disintegration, while less contain of polyacrylic acid would lead to bioactivity of the cement. In the present study we investigated the effect of BAG from the CaSiO_3_-Ca_3_(PO_4_)_2_ system addition to the bioactivity of RMGIC. The BAG containing 10, 15, and 20% of P_2_O_5_ (denoted as CSP10, 15, and 20) were used in the study to modify the powder of RMGIC, with apatite wollastonite (AWG) was chosen for a comparison. The surface bioactivity was assessed using XRD, FT-IR, and SEM analysis after specimen immersion in the simulated body fluid (SBF). Measurement of Ca, P, F, Sr, and Al was conducted for the remaining SBF. Cells studies were done to evaluate cell attachment, proliferation, and differentiation on the RMGIC containing BAG compared to the one without BAG. Results of Sr and Al analysis lead to the conclusion that addition of BAG may not influence stability of the matrix of the cement. It was also confirmed that addition of bioactive glass was positive factor indicating excellent ions exchange in SBF and spontaneous growth of apatite by consuming the Ca and P ions in the surrounding fluid. The ability of osteoblasts differentiation on the four types of bioactive cements were higher than that of RMGIC without BAG. These results might provide novel insights into the development of a new generation of osteoconductive biomedical materials.

## Introduction

Glass ionomer cements (GIC) are currently used for various dental applications. Over other restorative materials, GIC has advantages because it can be placed directly without an additional bonding agent into tooth cavities and has fluoride releasing property [1–4]. However, GIC also has some drawbacks related to its sensitivity to water at the initial stage of setting. At the initial stage of setting, GIC paste is potential to be exposed to saliva which decrease its mechanical strength significantly. The drawback of conventional GIC was then modified by water soluble resin [5] and called resin-modified glass ionomer cement (RMGIC). The basic composition of the liquid phase of RMGIC is polycarboxylic acid, 2-hydroxyethylmethacrylate (HEMA), and water. While the composition of the powder phase of RMGIC is basically the same as those of conventional GIC. The powder composition of the conventional GIC, that is fluoroaminosilicate glass, was designed to allow the glass to react with weak acid inside the glass system, that is poly-carboxylic acid [6].

Nowadays, the development of GIC is going further beyond dental material. It goes to the direction in which they are used for bone replacements and wider biomedical applications [7–12]. For that purpose, bioactivity is considered an important factor for the successful application. However, even if the powder phase of RMGIC contains both calcium and phosphate, so far RMGIC does not show any bioactivity. If modified GIC in the form of RMGIC could possess bioactivity, it would certainly be an advantage in the area of regenerative dentistry, but only a limited research has been done so far.

Matsuya and coworkers [10] reported a new GIC based on the composition of bioactive CaO-P_2_O_5_-SiO_2_(-MgO) glass. Using Fourie-transform infrared (FT-IR) and magic angle spinning-nuclear magnetic resonance (MAS-NMR) spectroscopies they investigated its setting related processes. Based on their investigation, carboxylate salt was formed due to released Ca^2+^ from bioactive glass (BAG). Besides, they also found that in the silicate network the degree of polymerization increased. The essential setting mechanism of the new cement through an acid base reaction between the basic glass and the polymeric acid was the same as the conventional GIC. However, compared to those of conventional GIC, the compressive strength of the set cement was much lower. The set cement also tended to disintegrate during SBF immersion. When they evaluated the bioactivity of the cement in the simulated body fluid (SBF), they found no apatite was formed on the surface of new set cement at least within 4 weeks.

Kamitakahara *et al*. also evaluated the possibility of obtaining bioactive GIC in the presence and absence of polyacrylic acid [13]. They analyzed apatite formation on the BAG when immersed in SBF. They found apatite formation was suppressed in the presence of 0.1 ppm of polyacrylic acid and GIC bioactivity was inhibited. It was known from the study of Kamitakahara and coworkers that polyacrylic acid suppresses the apatite forming ability of BAG.

In view of this, hybridization of RMGIC and BAG is one of the methods to overcome the limitations of the conventional GIC. This is because of less concentration of polyacrylic acid inside RMGIC compared to that of conventional GIC. Less concentration of polyacrylic acid is due to rapid set of RMGIC by HEMA or methacrylate groups polymerization in the polyacid chain. Modification of GIC by resin component inclusion in the cement was also expected in higher mechanical strength and resistance to the disintegration. Our previous study [14] showed that inclusion of four types of BAG into a commercially available RMGIC was clinically acceptable regarding their setting time, mechanical strength, and setting mechanism. Excellent setting ability was provided by the inclusion of BAG into RMGIC. Although Ana and coworkers [14] observed the decreased mechanical strength of the set cement containing BAG when compared with RMGIC, but it was much higher when compared to the conventional GIC containing BAG. Therefore, they concluded that it was possible to develop bioactive-RMGIC for clinical applications. From the study, it was recommended that the next step to be investigated is its bioactivity.

In this study we replaced the powder phase of a commercially available RMGIC with the BAG. The present study was aimed at examining the possibility of obtaining bioactive RMGIC wherein apatite forms of the cements, after immersion in the SBF, were investigated by X-ray diffraction patterns, FT-IR spectroscopies, chemical analysis and microstructural changes. Cell studies were also done to predict its biocompatibility and bioactivity.

## Materials and Methods

### Preparation of bioactive powders

Four BAG powders were prepared and used in this study, referred to the previous study [14] as depicted in **Table 1**. Glass CaSiO_3_-Ca_3_(PO_4_)_2_ system was used to compose BAG powders to achieve 10, 15, and 20 weight% P_2_O_5_ content which was then respectively encoded as CSP10, CSP15, and CSP20. For the comparison, formulation of Apatite Wollastonite (AW) glass-ceramics as reported by Kokubo [12] was also prepared without adding CaF2 and encoded as AWG in the following text.

**Table 1.** Composition of BAG prepared in the study (in mol%) as referred to the previous study [14].

| Glass | CaO | SiO<sub>2</sub> | P<sub>2</sub>O<sub>5</sub> | MgO | Remarks |
| --- | --- | --- | --- | --- | --- |
| CSP10 | 54.3 | 41.3 | 4.3 | - | CaSiO<sub>3</sub>-Ca<sub>3</sub>(PO<sub>4</sub>)<sub>2</sub><br>10wt% as P<sub>2</sub>O<sub>5</sub> |
| CSP15 | 56.7 | 36.6 | 6.7 | - | CaSiO<sub>3</sub>-Ca<sub>3</sub>(PO<sub>4</sub>)<sub>2</sub><br>15wt% as P<sub>2</sub>O<sub>5</sub> |
| CSP20 | 59.2 | 31.6 | 9.2 | - | CaSiO<sub>3</sub>-Ca<sub>3</sub>(PO<sub>4</sub>)<sub>2</sub><br>20wt% as P<sub>2</sub>O<sub>5</sub> |
| AWG | 50.1 | 35.6 | 7.2 | 7.1 | - |

### Preparation of RMGIC cement containing BAG

Resin- modified glass ionomer cement containing BAG was prepared by addition of BAG into the powder of RMGIC (Fuji II LC, GC Corp., Tokyo, Japan), denoted as FLC in the following text. The detailed composition of Fuji II LC powder phase is not publicly open, but it is reported that the powder is free from polycarboxylic acid. The powder mixture of FLC and BAG were mixed to achieve 33 or 50 weight% BAG contents. Notation of the prepared cement powders are described as BAG added and its content in parentheses, as follows: CSP10(33), CSP10(50), CSP15(33), CSP15(50, CSP20(33), CSP20(50), AWG(33) and AWG(50). Without any modification, Fuji II LC liquid was used and applied as the liquid phase of BAG containing cement for simplicity. The cement powder and liquid were mixed and poured into Teflon molds in 6 mm diameter by 0.7 mm thickness for bioactivity test in SBF, 10 mm diameter by 0.1 mm thickness for cell attachment test, and 6 mm diameter by 0.7 mm thickness for cell proliferation and differentiation test). Light irradiation was done for 2X20 seconds after 2 minutes from the start of mixing. Specimens were then incubated at 37° C and 100% humidity for 24 hours to allow it to set completely. All the steps in preparing RMGIC containing BAG were referred to previous study [14].

### Immersion in SBF

Disc specimens were soaked in 20 mL SBF at 37° C for various periods to test bioactivity in vitro. The SBF was prepared to reach human blood plasma ion concentrations at 7.4 pH level. The ion concentrations of the SBF solutions used in this study was referred to the one given by Kokubo [15] as described in **Table 2**. After 0, 1, 2, 4, and 8 weeks, the specimens were washed with distilled water, dried, and prepared for analysis.

**Table 2.** Chemical composition of SBF [15].

| Order | Reagent | Amount (g/L) |
| --- | --- | --- |
| 1. | NaCl | 7.996 |
| 2. | NaHCO<sub>3</sub> | 0.350 |
| 3. | KCl | 0.224 |
| 4. | K<sub>2</sub>HPO<sub>4</sub>.3H<sub>2</sub>O | 0.228 |
| 5. | MgCl<sub>2</sub>.6H<sub>2</sub>O | 0.305 |
| 6. | 1 mol/L HCl | 40cm<sup>3</sup> |
| 7. | CaCl<sub>2</sub> | 0.278 |
| 8. | Na<sub>2</sub>SO<sub>4</sub> | 0.071 |
| 9. | (CH<sub>2</sub>OH)<sub>3</sub>CNH<sub>2</sub> | 6.057 |
| 10. | 1 mol/L HCl | Adjust pH by dropping |

### Surface analysis after immersion

The specimens were studied by means of X-ray diffraction (XRD) to confirm whether apatite was present after soaked in the SBF for various periods at typical scan step 0.02° 2*θ* and scan speed 2°/min. A Cu Kα tube was operated at 40 kV and 100 mA was used for the generation of X-ray (RINT 2500V, Rigaku Co., Tokyo, Japan). The surface of specimens were examined by type JSM 5400 LV (JEOL, Tokyo, Japan) scanning electron microscope (SEM) after gold sputtering on the surface of the specimens using JEC 550 Twin Coater (JEOL, Tokyo, Japan) to reconfirm whether apatite was formed on the surface of RMGIC containing BAG. To assess type of apatite formation on the surface of specimens, Fourie-transform infrared (FT-IR) was applied to the ground surface. The transmission spectra of FT-IR were obtained using an IR spectrometer (Spectrum 2000, The Perkin-Elmer Co., CT, USA) by KBr disc method.

### Chemical analysis of immersion solution

The SBF used for immersion was chemically analyzed for pH, fluoride, aluminum, strontium, calcium, and phosphate before and after the specimen’s immersion. Level of pH was determined using glass pH meter (TPX-90, Toko Chemical Lab. Co. Ltd., Tokyo, Japan). Fluoride concentration was analyzed by an ion specific combination electrode (model 720A plus, Orion research, Cambridge, MA). To measure the fluoride concentration of each sample at a specific time, 2 mL of each were placed in a plastic cup. In order to adjust ionic strength and pH, the sample was mixed with 2 mL of 10 times diluted TISAB III solution (Orion research, Cambridge, MA). The fluoride electrode was calibrated just before each measurement using the same solution mixed with 2 mL of solutions containing fluoride 0.1, 0.2, 0.5, 1, and 5 ppm. Although reproducibility was not strictly checked, but remeasurement was conducted to always give less than 10% concentration difference.

Calcium (Ca 422.7nm), aluminum (Al 309.3 nm), and strontium (Sr 460.7nm) concentrations were determined by atomic absorption spectroscopy (Analyst 300, The Perkin-Elmer Co., Tokyo, Japan) with C_2_H_2_/N_2_O, C_2_H_2_/N_2_O, and C_2_H_2_/ air flame respectively. To suppress ionization interference for the Ca analysis after immersion, 0.5 mL SBF solution was mixed with of 1 mol/L hydrochloric acid and 1 mL of 1% KCl. The SBF solution without Ca element was also prepared as standard solution with 2.0 to 5.0 ppm for Ca analysis. Meanwhile, in order to analyze Al and Sr by suppressing ionization interference, 5 mL of each sample was mixed with 1 mL of 1 mol/L hydrochloric acid and 1 mL of 1% KCl. Standard solutions for Al and Sr element were between 0.1 to 5.0 ppm and for Ca element were between 2.0 to 5 ppm. Each measurement was repeated in triplicate and the concentration values were averaged. Standard deviation was always set up less than 5%. The amount of Ca concentration from each specimen was expressed as ppm. The amount of each Al and Sr species released from the specimen was expressed as a cumulative amount per unit surface area of specimen disc (μg/cm^2^).

Phosphate concentration of each SBF solution after immersion was determined using ultraviolet/ visible light spectrometer (UV/ VIS Spectrometer, U-best 50, JASCO, Tokyo, Japan) in which 1 mL of each sample was mixed with 1 mL Na_2_MoO_4_.2H_2_O 2.5 gram/L and 1 mL 0.5 N_2_H_6_SO_4_. Standard solution of P was prepared from KH_2_PO_4_ with concentrations between 0.5 to 2.0 ppm. Each measurement was repeated five times and the concentration values were averaged. The amount of P concentration from each specimen was expressed as ppm.

### Cell attachment, proliferation, and differentiation

Osteoblast like cells (MC3T3E1) was used in the study to confirm bioactivity. Cells were obtained from Gibco (USA). The culture medium was α-MEM (Eagle, Sigma Chemical Co., St. Louis, MO, USA) with 10% fetal bovine serum (FBS, Gibco, Invitrogen Co., Auckland, New Zealand) in an atmospheric conditioned containing 5% CO_2_ at 37° C. Cells were seeded on each disc with 1X10^4^ cells concentrations per 10 mm disc and plastic dishes (24 well cell culture cluster, Corning, New York, USA) containing no osteoblasts were used as the control.

Onto each specimens, the cells were seeded and incubated for 5 hours. To eliminate non-adherent cells, the specimens were washed with 0.1 mol/L phosphate buffer saline (PBS) after 5 hours incubation. Report for cell attachment was conducted by harvesting the adherent osteoblasts on the specimens, continued by counting using Burker and Turk hemocytometer. The PBS containing collagenase and trypsin was prepared for adherent cells harvesting.

Cell proliferation was measured at 1, 3, 5, 7, and 9 days using MTT assay. In brief, the reagent, MTT (3[4,5-dimethylthiazole-2-yl]2,5-diphenyltetrazolium bromide) was enzymatically counted by living cells as formazan product. In vitro cells proliferation was directly related to the color intensity of the produced formazan which expressed the number of viable cells. MTT reagent was added to each sample and incubated at 37° C for 4 hours. Dimethyl sulfoxide was used to solubilize formazan product. Absorbance measurement was conducted at 570 nm using a plate reader (MTP-32 Microplate Reader, Corona Elect., Ibaragi, Japan).

Alkaline phosphatase (ALP) activity of osteoblasts on day 6, 9, and 12 was determined by enzyme-linked immunosorbent assay (ELISA) using p-nitrophenyl phosphate as the substrate (ALP substrate system, Kierkegaard and Perry Lab., Gaithersburg, MD, USA). The absorbance of p-nitrophenol at 405 nm was determined with plate reader (MTP-32 Microplate Reader, Corona Elect., Ibaragi, Japan). Enzymatic activity was expressed as nmol of p-nitrophenol/ minute per mg protein. The protein content was determined by the method of Bradford (1976) using bovine-γ-globulin as a standard [16].

For type I procollagen peptide (PICP) and osteocalcin analysis, each specimen was decalcified with 0.6 mmol/L HCl for 30 minutes. The obtained solution was harvested and stored at −20°C until assay. The amount of PICP on day 3, 9, and 12 and osteocalcin on day 9, 15, and 21 were measured with an enzyme immunosorbent assay (ELISA) kit (Takara, Shiga, Japan).

### Statistical analysis

For the statistical analysis, one-way ANOVA and Fischer’s PLSD method as post-hoc-test were performed. A p value of <0.05 was considered statistically significant.

## Results

### Surface analysis on the bioactivity of the set cement after immersion in SBF

Bioactivity could be evaluated by the apatite formation on the surface of the cement after immersion in SBF. **Figures 1** and **2** show respectively X-ray diffraction patterns of FLC (RMGIC without BAG), as a control, and the cement containing 50 weight% of CSP10, CSP15, CSP20 and AWG after various immersion time in SBF, with hydroxyapatite (HAP) diffraction pattern as a reference. With the control, FLC, no apatite peak was observed at any period of immersion as shown in **Figure 1**. Diffraction patterns for the cement containing BAG show two broad diffraction peaks around 25.5 and 32.0° assigned to apatite phase [16–17] after 4 or 8 weeks immersion on the surface of CSP10(50), CSP15(50), and CSP20(50), as well as 8 weeks after immersion on AWG(50). The study also confirmed that apatite formation was also observed in the cements containing 33% of BAG except for AWG. However, the formation was delayed and confirmed only after 8 weeks immersion in SBF.

**Figure 1.**
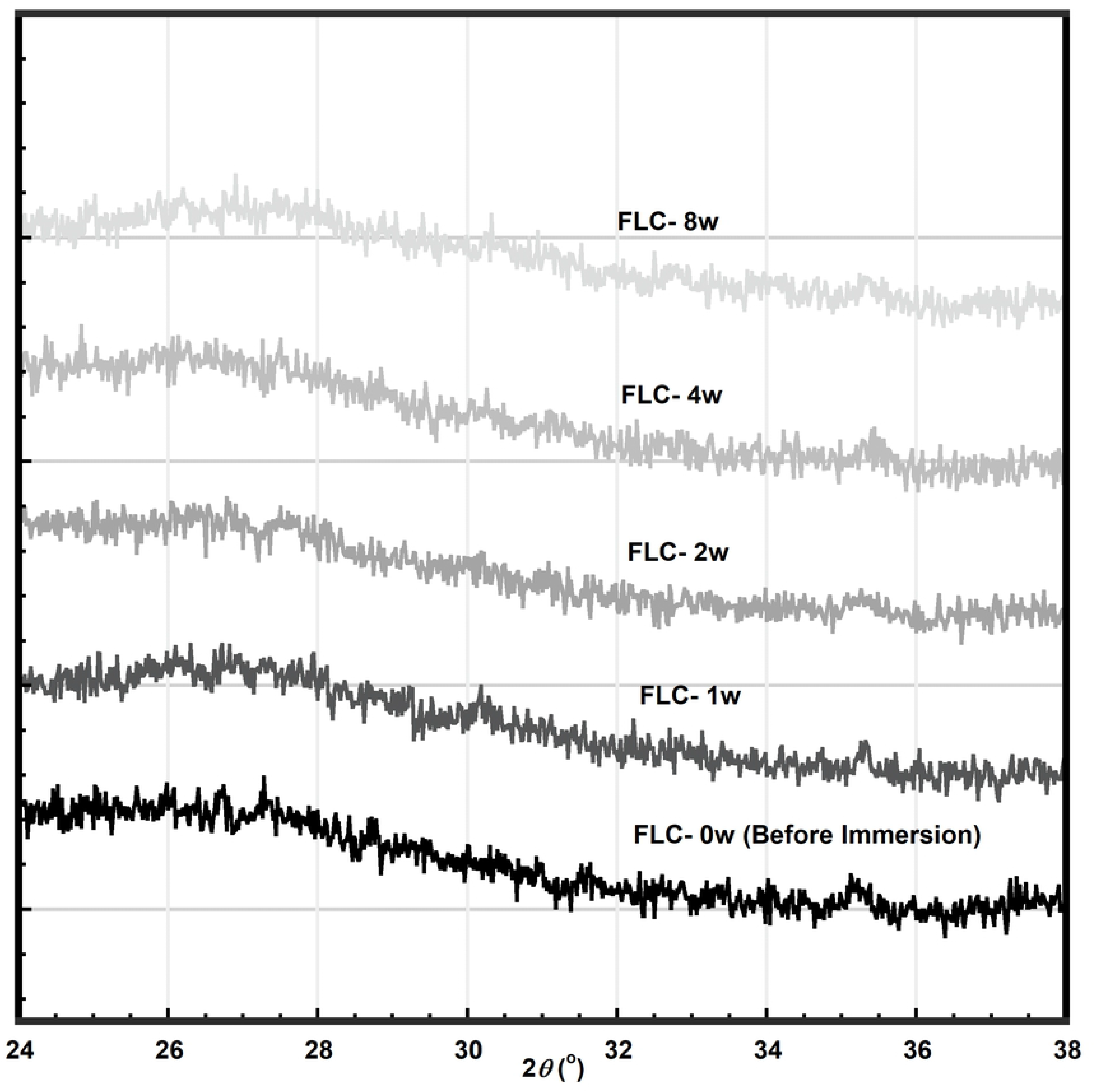
X-ray diffraction pattern of resin modified glass ionomer cement (FLC) after immersion in SBF at various time up to 8 weeks.

**Figure 2.**
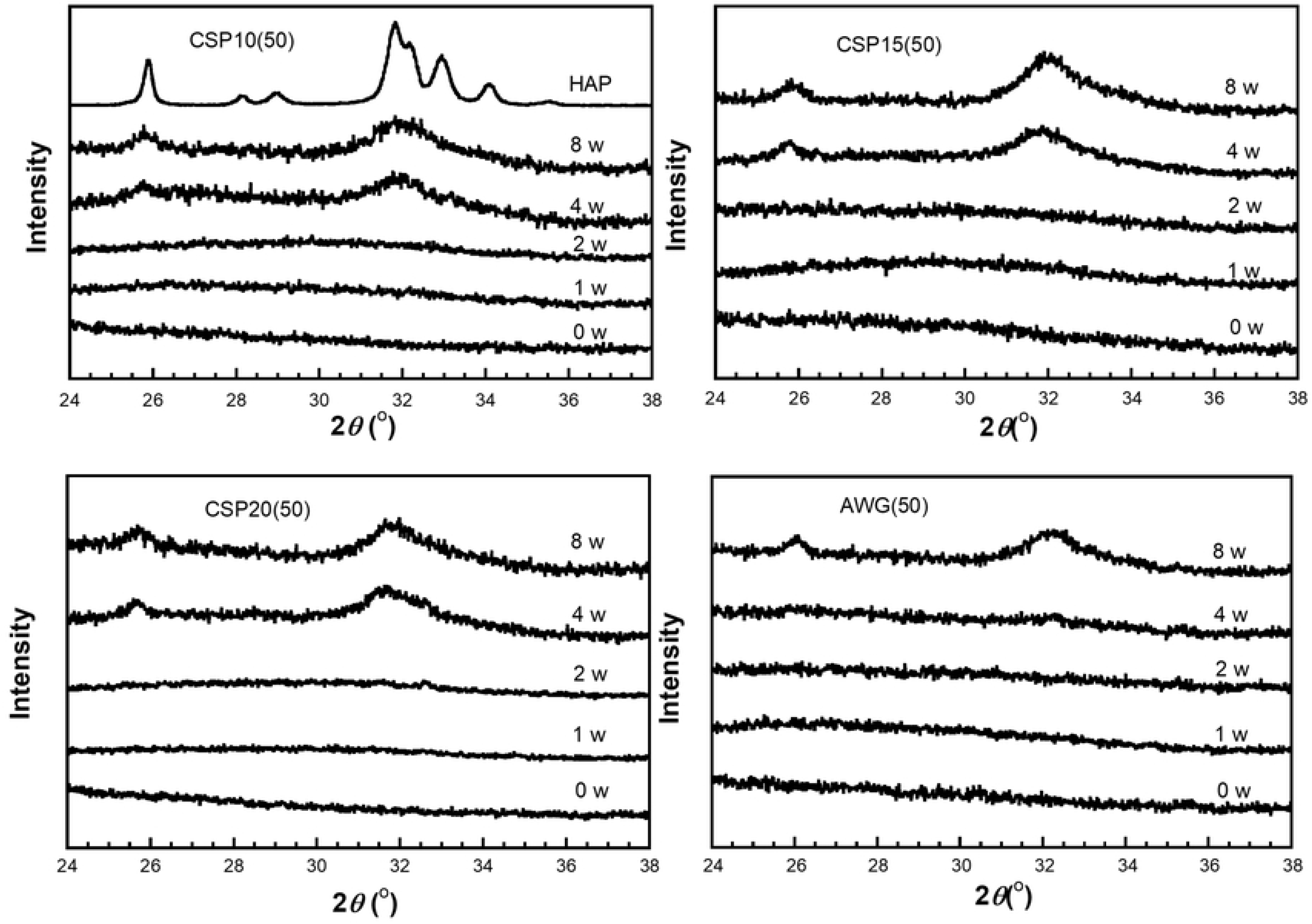
X-ray diffraction pattern of resin modified glass ionomer cement containing bioactive glass CSP10(50), CSP15(50), CSP20(50), and AWG after immersion in SBF at various time up to 8 weeks. HAP shows diffraction of synthetic hydroxyapatite.

Bioactivity of the cement containing BAG was also detected by microstructure of the specimens observed with SEM. **Figure 3** shows representative of the micrograph of the specimens surface observed in this study, those are FLC, CSP10 in 50 weight%, and AWG in 50 weight%. As confirmed from the XRD patterns shown in **Figure 1** and **2**, Figure 3 confirmed spherical crystalline apatite layer was formed on the surface of the cements after immersion for 4 and/or 8 weeks. With FLC, any crystalline materials were not observed even after immersion for 8 weeks, as expected from the XRD result.

**Figure 3.**
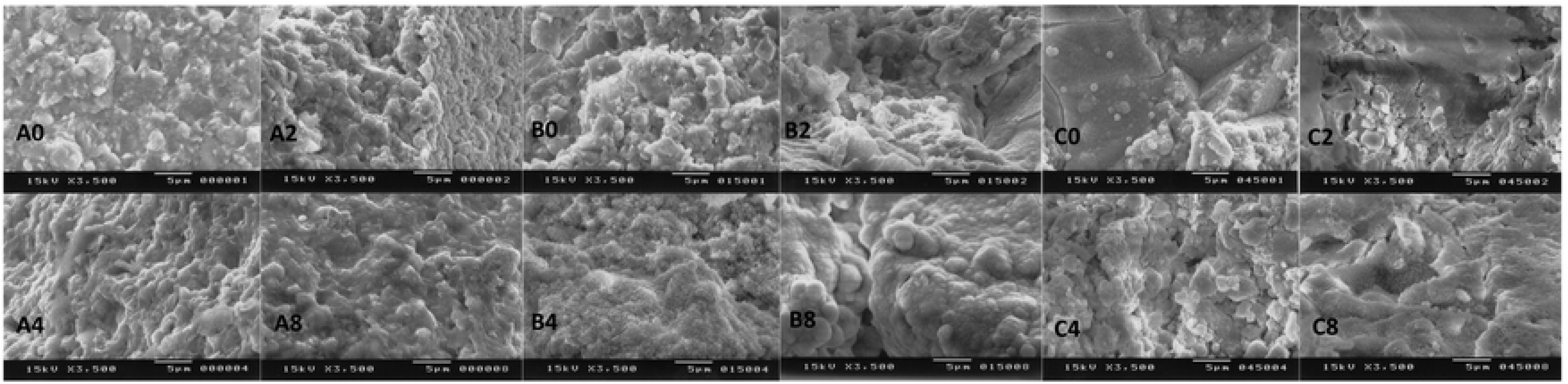
Scanning electron micrograph representatives of (A) resin modified glass ionomer cement (FLC), (B) CSP10(50), and (C) AWG after 0, 2, 4, and 8 weeks immersion in SBF.

The FT-IR spectra, as shown on **Figure 4**, detected that there was no change on the spectra of FLC before and after immersion, except for an increase in intensity of a peak at 1563 cm^-1^ assigned to asymmetric vibration of carboxylate. On the other hands, the FT-IR spectra for CSP10(50) showed new absorption bands around 873, 602 and 568 cm^-1^ assigned to CO_3_ and PO_4_ in carbonated apatite besides the asymmetric stretching vibration of the carboxylates at 1563 cm^-1^. The bands at 1460 and 1415 cm^-1^ assigned to CO3 in B-type carbonated apatite overlapped with those from symmetric vibration mode of carboxyl group and ester in the original cement, FLC. Increase in intensity of the absorption bands, however, clearly showed the formation of carbonated apatite after immersion. Other cements, CSP15(50), CSP20(50) and AWG(50) also revealed similar change in FT-IR spectra as shown in **Figure 5** together with CSP10(50) and FLC. The same indication on the bioactivity of the cements was also found in CSP10(33), CSP15(33), and CSP20(33) after 4 weeks immersion in SBF. In general, the bioactivity of the cements containing BAG after immersion in SBF was confirmed by XRD, SEM, and FT-IR as concluded in **Table 3**.

**Figure 4.**
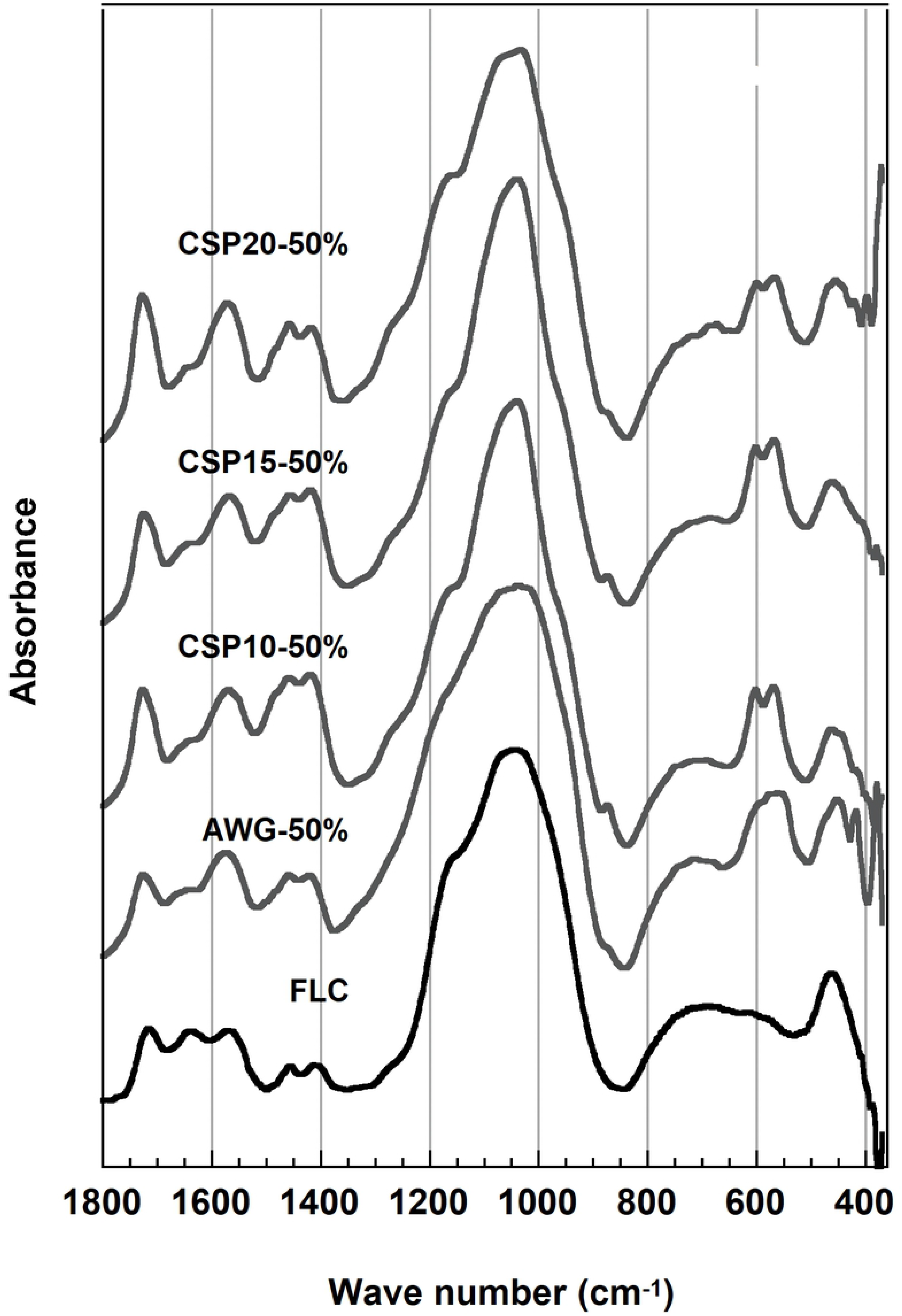
FT-IR spectra of the cements containing 50% of bioactive glasses and FLC after 8 weeks immersion in SBF.

**Figure 5.**
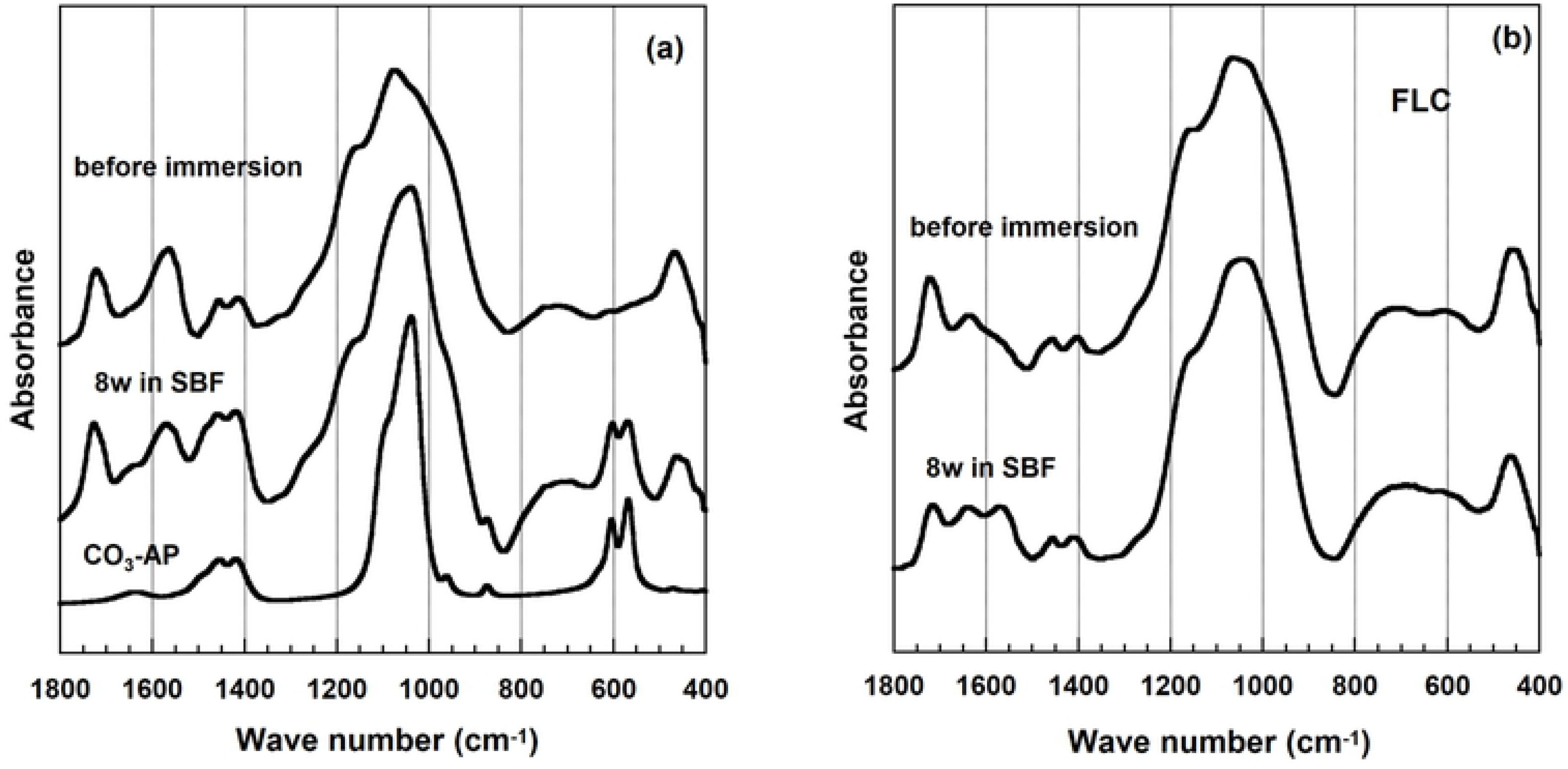
FT-IR spectra for (a) resin modified glass ionomer cement containing bioactive glass CSP10(50) and (b) FLC before and after 8 weeks immersion in SBF. Spectra of carbonated hydroxyapatite (CO3-Ap) are used as a reference.

**Table 3.** Apatite formation in the cements after immersion in SBF 37°C.

| Content<br>(weight %) | Time<br>(week) | Formation of Apatite on the Cement<br>After Immersion in SBF<br>(XRD, FTIR, and SEM Confirmation) |  |  |  |  |
| --- | --- | --- | --- | --- | --- | --- |
|  |  | FLC | CSP10 | CSP15 | CSP20 | AWG |
| 33 | 1 | - | - | - | - | - |
|  | 2 | - | - | - | - | - |
|  | 4 | - | - | - | - | - |
|  | 8 | - | Yes | Yes | Yes | - |
| 50 | 1 | - | - | - | - | - |
|  | 2 | - | - | - | - | - |
|  | 4 | - | Yes | Yes | Yes | Yes |
|  | 8 | - | Yes | Yes | Yes | Yes |

### Calcium and phosphate concentration in SBF solution after immersion

**Figures 6** shows Ca^2+^ ions concentration in SBF after certain periods of immersions. It was found that calcium concentration tended to increase by the addition of BAG. This tendency was not found in FLC, e.g. calcium concentration tended to decrease in FLC. Meanwhile, phosphate concentration analysis shows opposite results as it is shown on **Figure 7**. The concentration of phosphate tended to decrease by addition of BAG and to increase in case of FLC. To give additional information, no significant changes of pH level during this study. The level of pH ranged from 7.4 – 8.1.

**Figure 6.**
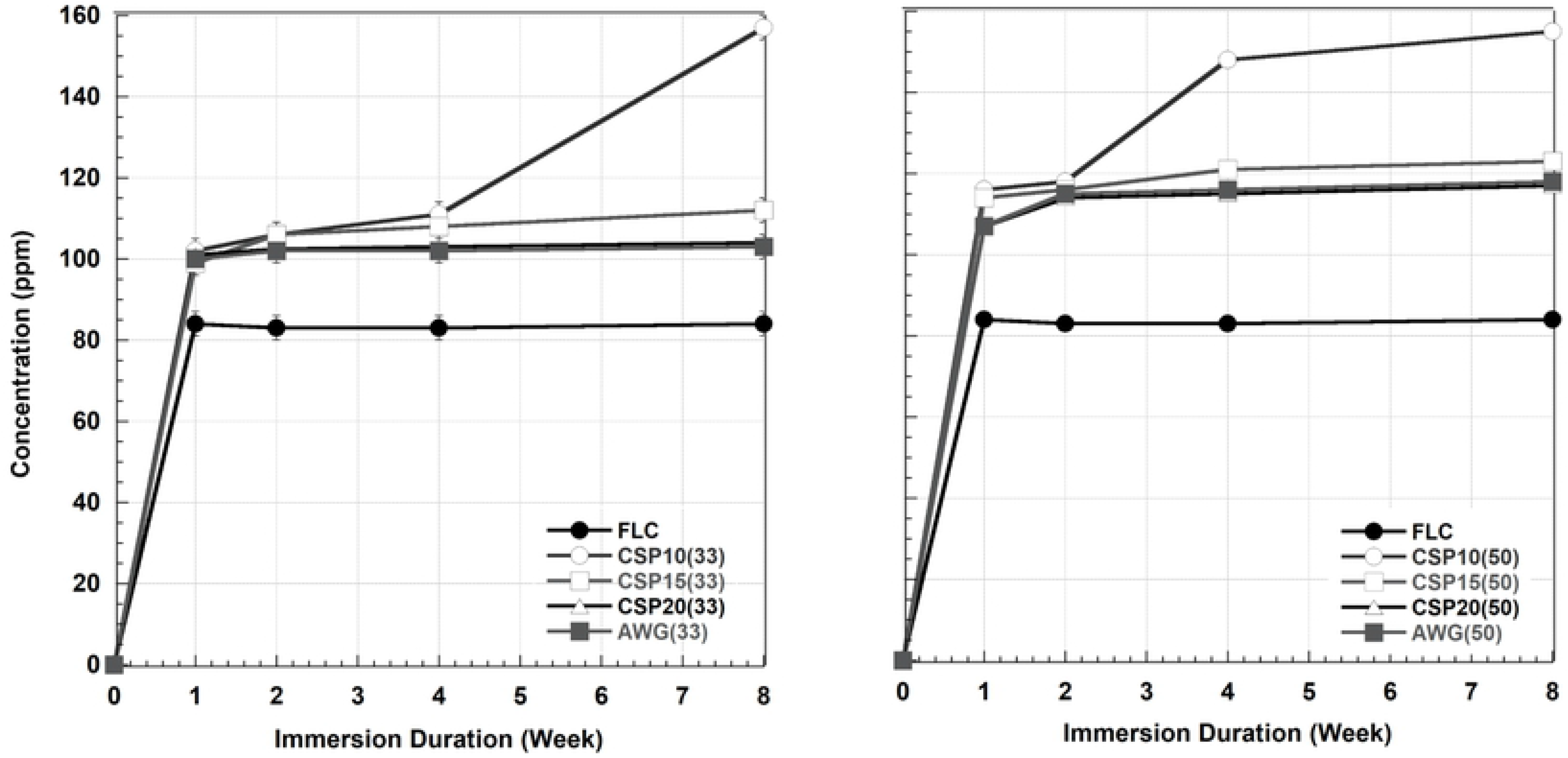
Calcium concentration in the remaining SBF after immersion of the cement without BAG (FLC) and with 4 types of BAG in 33 and 50% for various immersion times.

**Figure 7.**
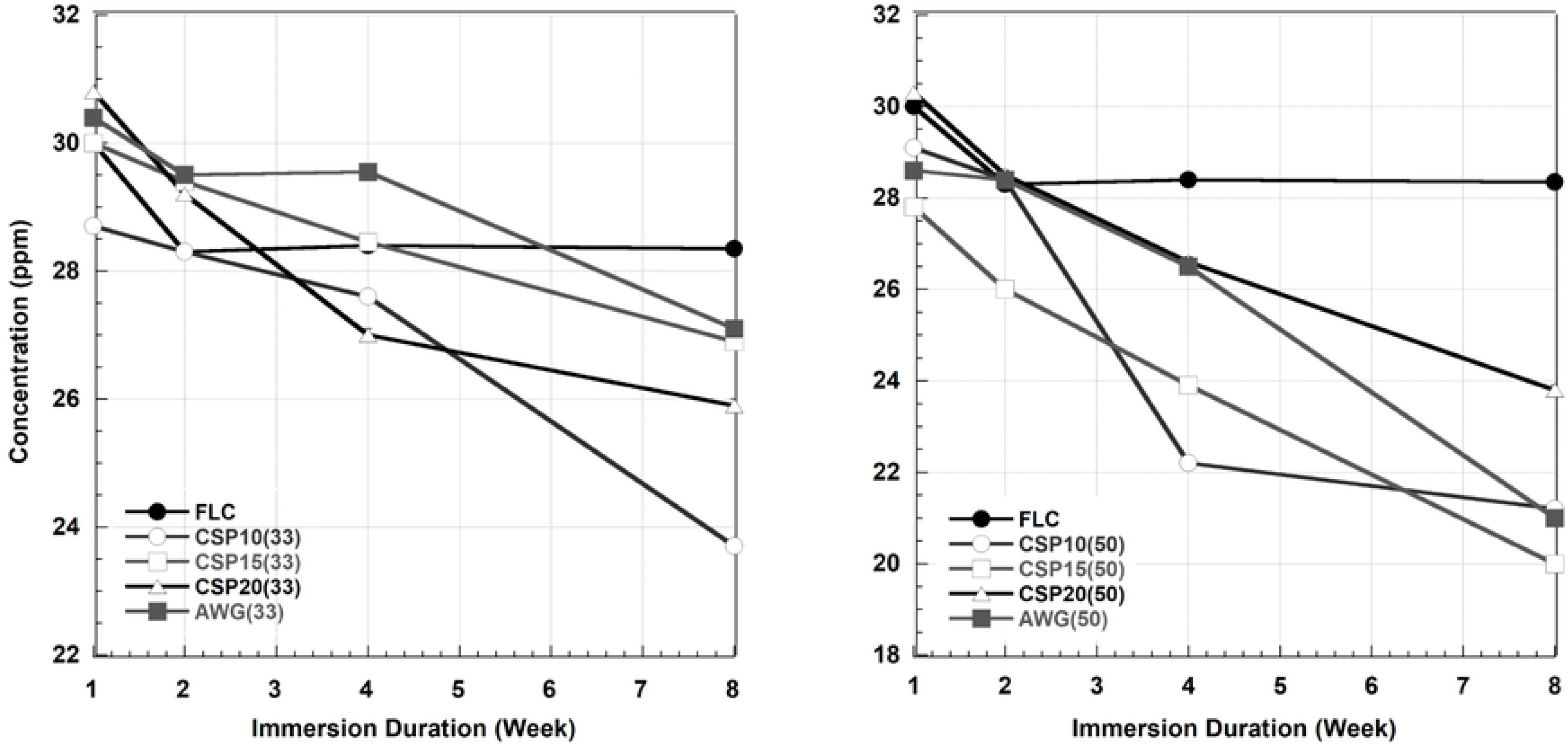
Phosphate concentration in the remaining SBF after immersion of the cement without BAG (FLC) and with 4 types BAG in 33 and 50% for various immersion times.

### F, Sr, and Al releases from the cement after immersion in SBF

**Figures 8** shows log-log plots for fluoride released from the various cements containing 33% and 50% BAG after immersion in SBF. It was found that F released into SBF decreased by glass addition and the extent of the decrease was larger in the cement containing 50% BAG. The finding was quite reasonable because the added BAG does not contain any fluoride ions so that fluoride release must have originated entirely from the powder component of the RMGIC. The amount of fluoride releases was also different among the types of the bioactive glasses and the order was different depending on the amount of addition. Although fluoride released from RMGIC containing BAG was lower than FLC, it was found that the bioactive RMGIC presented a pattern of fluoride release parallel to the control or reference (FLC).

**Figure 8.**
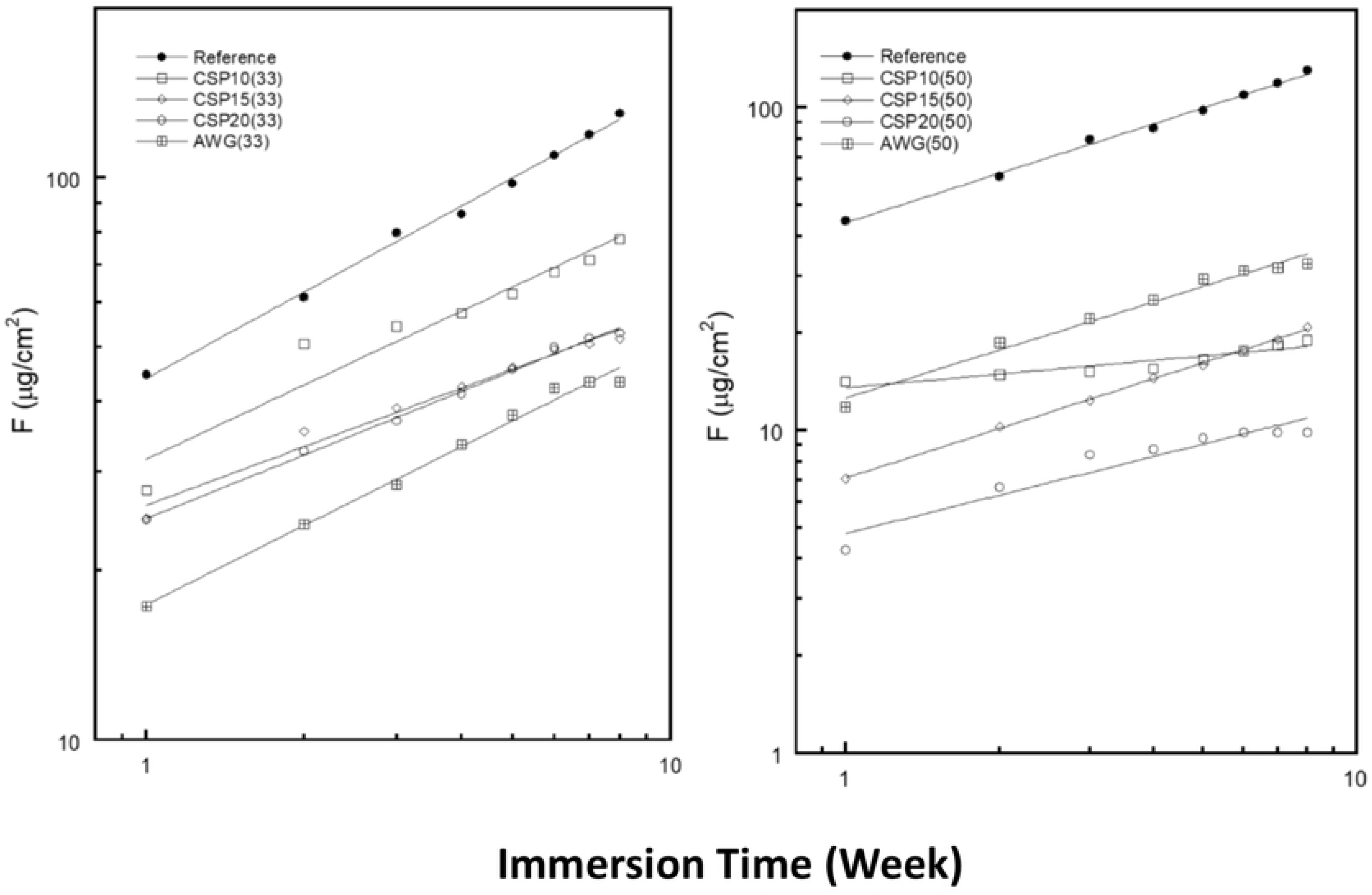
Log-log plot for fluoride (F) released in SBF from the reference (FLC without BAG) and cement containing 4 types BAG in 33 and 50% after various immersion times.

**Figure 9** shows log-log plots of Sr released. It was also found that the amount of Sr released from the RMGIC decreased by the addition of BAG as observed in the fluoride release. Unlike the F and Sr releases, the amount of Al released increased by the addition of the BAG, though the total amount of Al in the cement decreased with the addition, as shown in the log-log plots in **Figure 10**.

**Figure 9.**
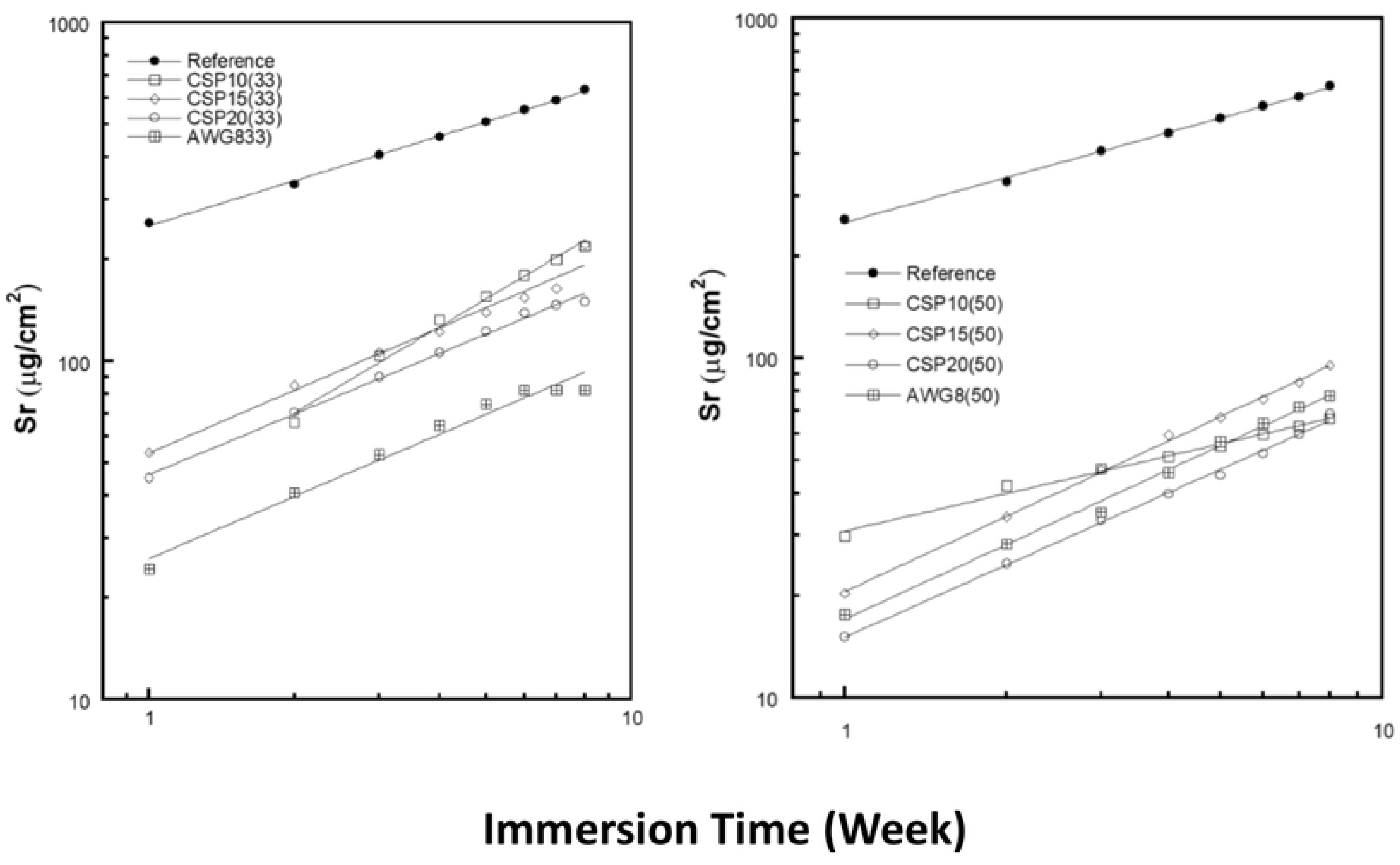
Log-log plot for strontium (Sr) released in SBF from the reference (FLC without BAG) and cement containing 4 types BAG in 33 and 50% after various immersion times.

**Figure 10.**
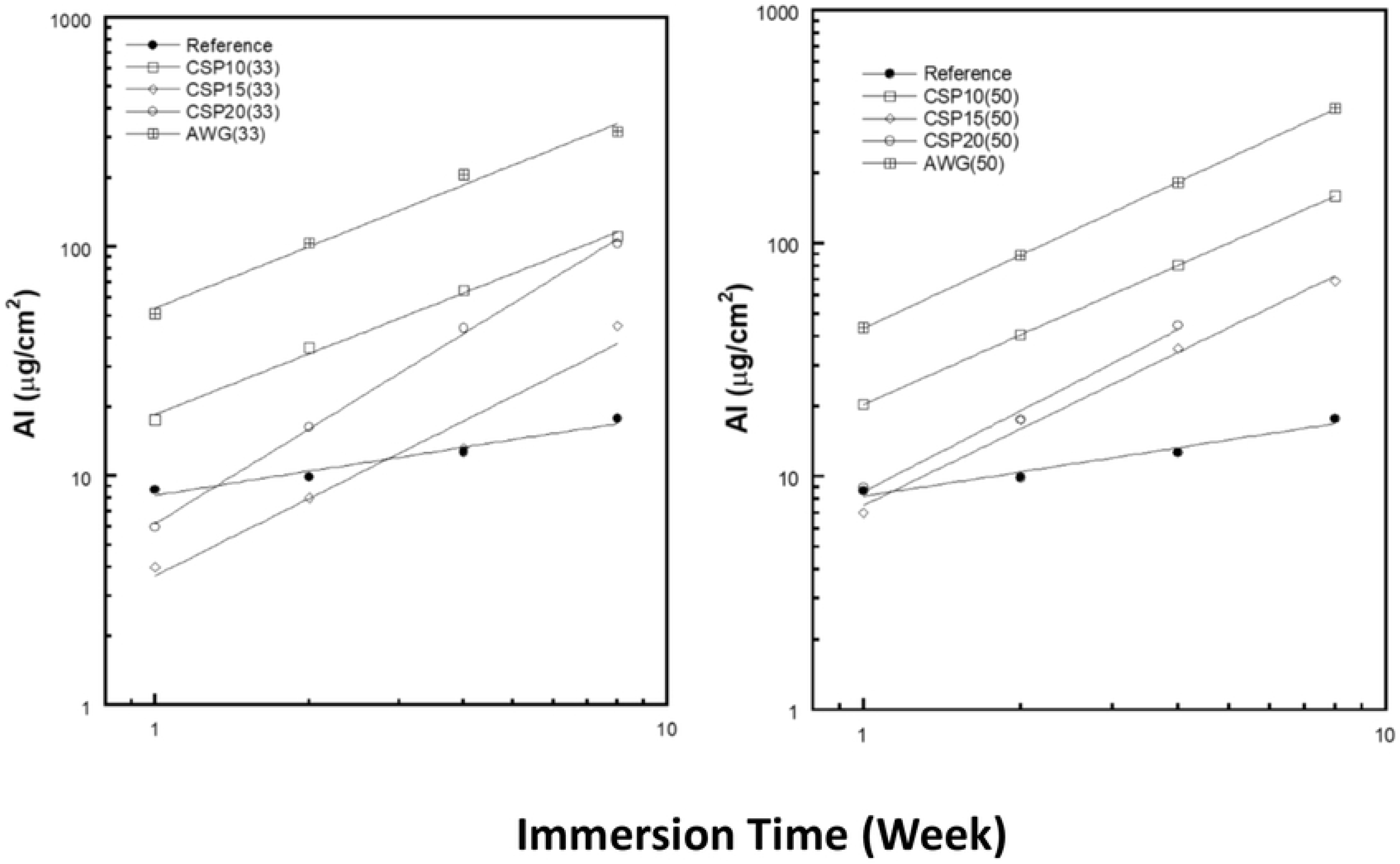
Log-log plot for aluminum (Al) released in SBF from the reference (FLC without BAG) and cement containing 4 types BAG in 33 and 50% after various immersion times.

### Attachment, proliferation, and differentiation of the cells on the cement

After 5 hours incubation initially, the cells attached on the surface of RMGIC containing BAG was significantly larger (p<0.05) compared to FLC (**Figure 11**). Regarding the type of the BAG, significant differences were found on the surface of RMGIC containing AWG glass with other types of resin- modified glass ionomer cement containing bioactive glass. As shown in **Figure 12**, the proliferation of MC3T3E1 cells on each sample increased correlatively with incubation time, and there were no significant differences among the four types of RMGIC containing bioactive glasses and FLC.

**Figure 11.**
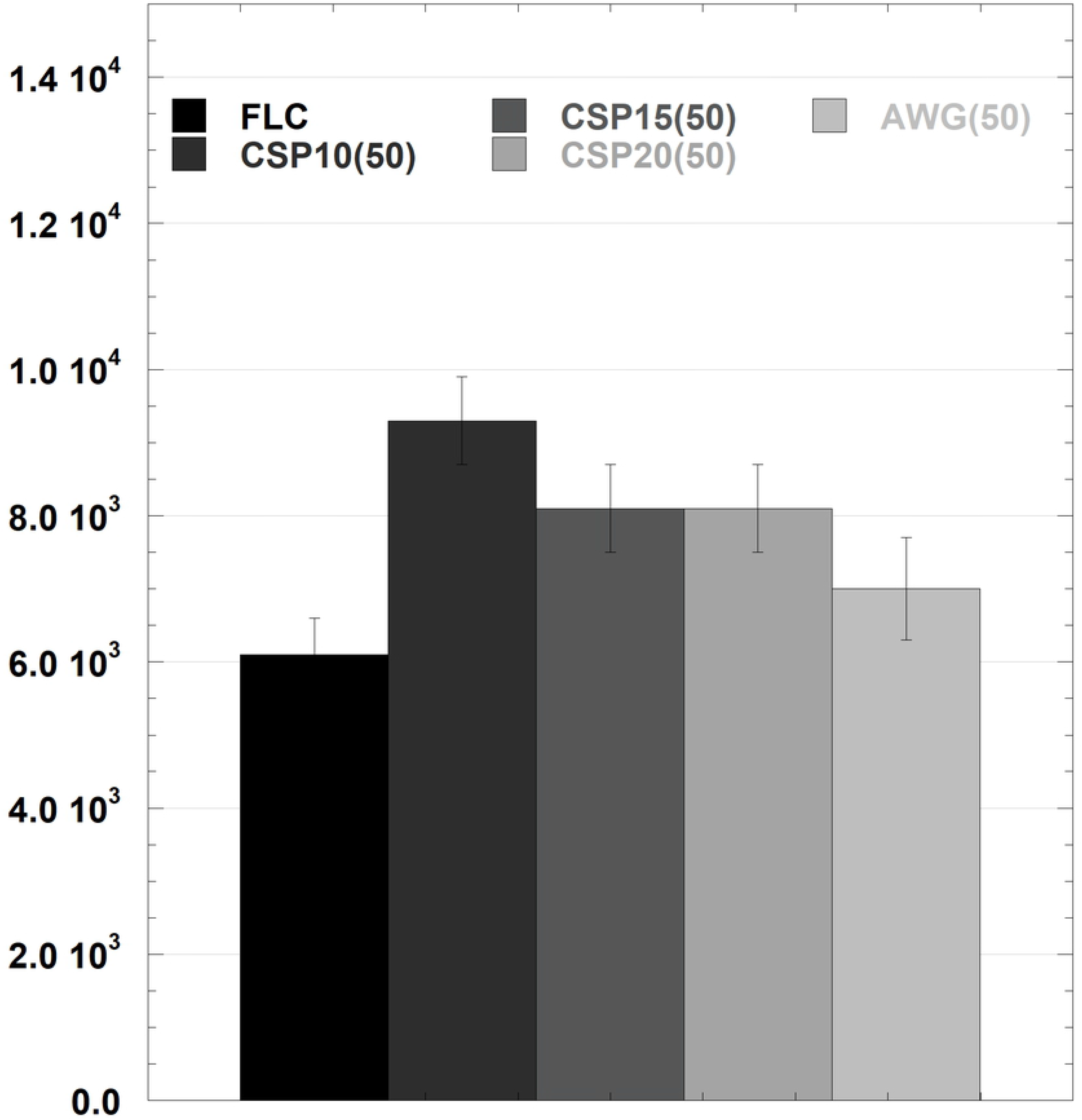
Cell attachment on the surface of FLC and cement containing 4 types BAG after 5 hours culture (n=3). The results are shown as the mean + standard deviation of the cells attached on the cement discs.

**Figure 12.**
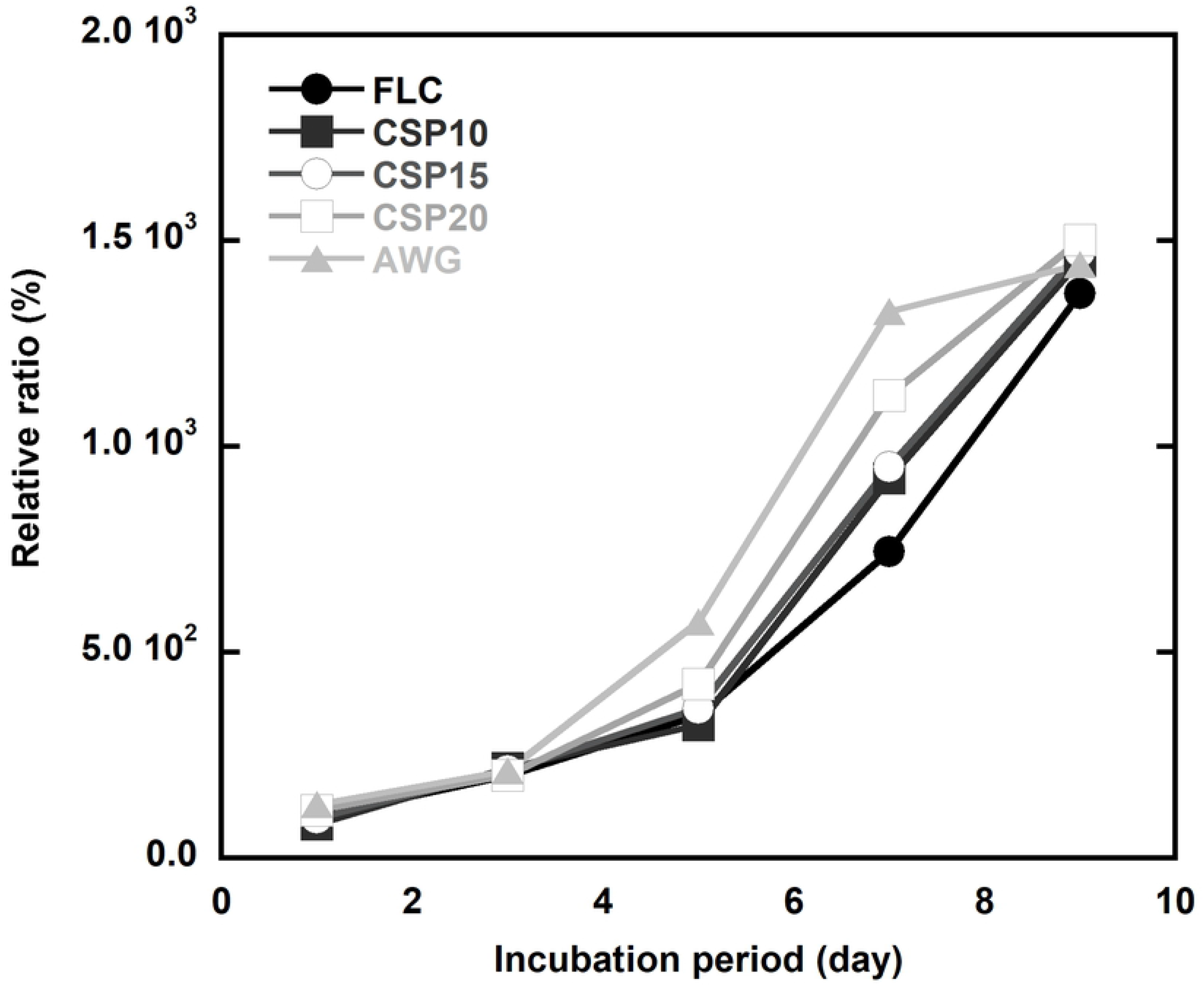
Osteoblast like cells proliferated on the surface of the FLC and cement containing 4 types BAG discs, evaluated by MTT assay. Each value was expressed as a percentage of the amount of absorbance produced by the cells on day 2. The results are shown as the mean + standard deviation (n=9).

**Figure 13** summarizes the course of type I collagen mRNA of osteoblasts on each specimen. The marked expression of type I collagen mRNA for all specimens increased with the incubation time. The cement containing BAG showed higher differentiation profile compared to FLC on the same day. For ALP mRNA, the same phenomena were found with the one of type I collagen mRNA (**Figure 14**). On day 12 AWG and CSP10 showed highest ALP mRNA value, CSP15 and CSP20 showed intermediate value, and FLC showed the lowest one.

**Figure 13.**
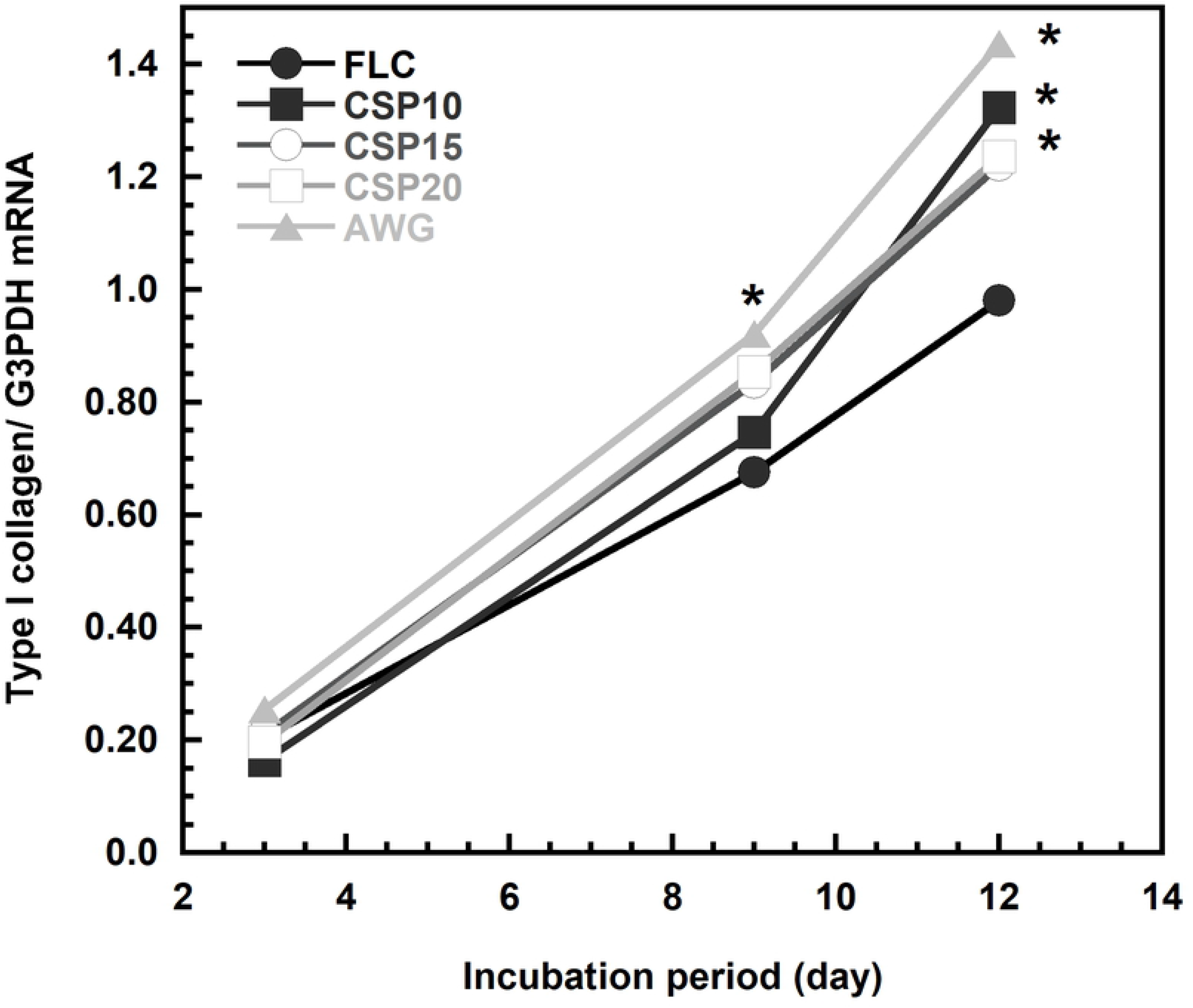
Time course of type I collagen mRNA expressed by MC3T3E1 on each cement specimens. The results are shown as the mean + standard deviation (n=9). The superscription indicates a significant difference as compared with FLC on the same day.

**Figure 14.**
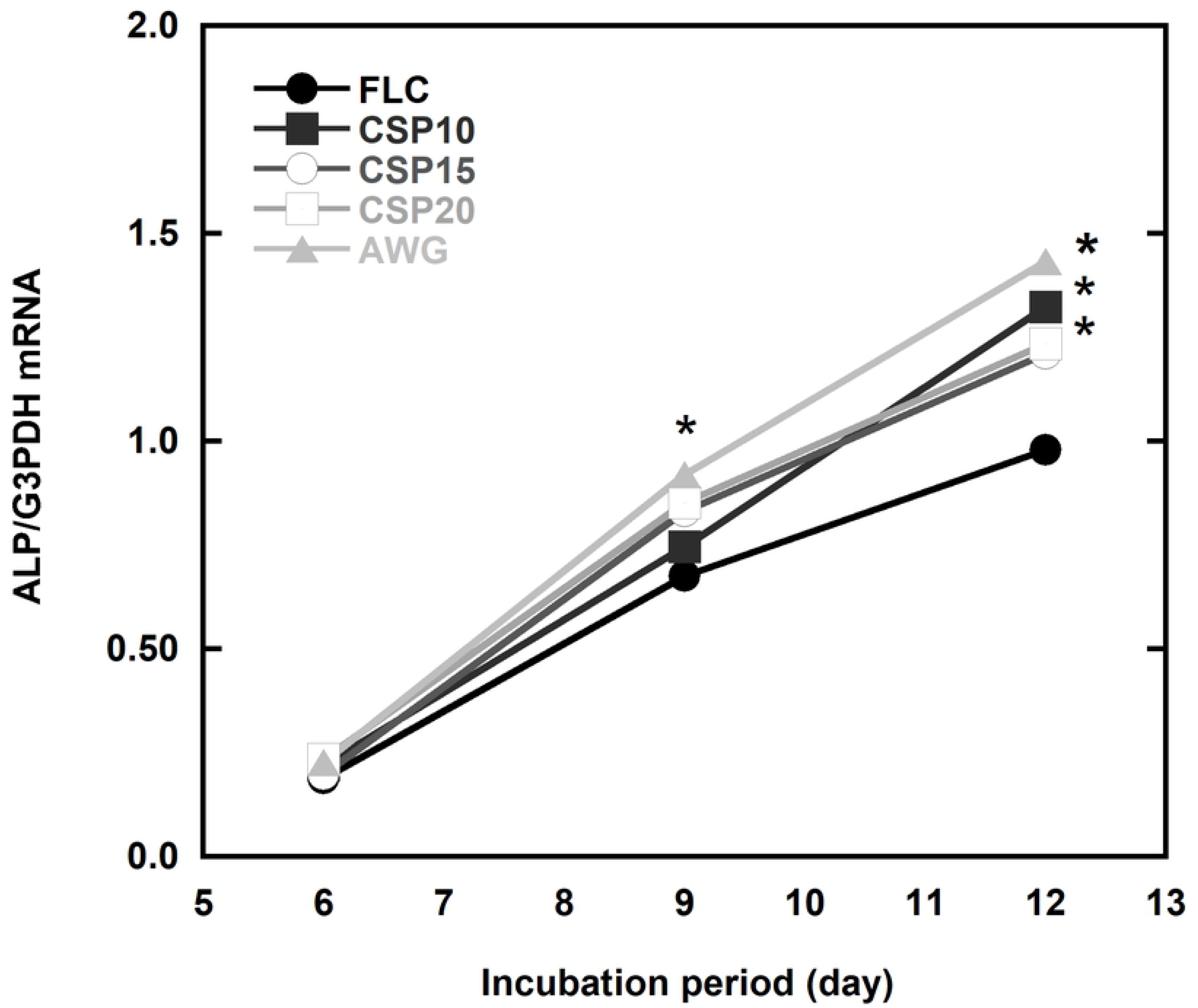
Time course of ALP mRNA expressed by MC3T3E1 on each cement specimens. The results are shown as the mean + standard deviation (n=9). The superscription indicates a significant difference as compared with FLC on the same day.

The gene expression of osteocalcin mRNA in each specimen (**Figure 15**) increased with the incubation time. Among all specimens, the increase rate of osteocalcin on AWG was the largest and in contrast FLC showed the slowest increase. As a result, AWG showed significantly (p<0.01) the highest value on day 15 and 21. Other resin-modified glass ionomer cement containing CSP10, CSP15, and CSP20 also showed higher differentiation profile compared to FLC, especially on day 21.

**Figure 15.**
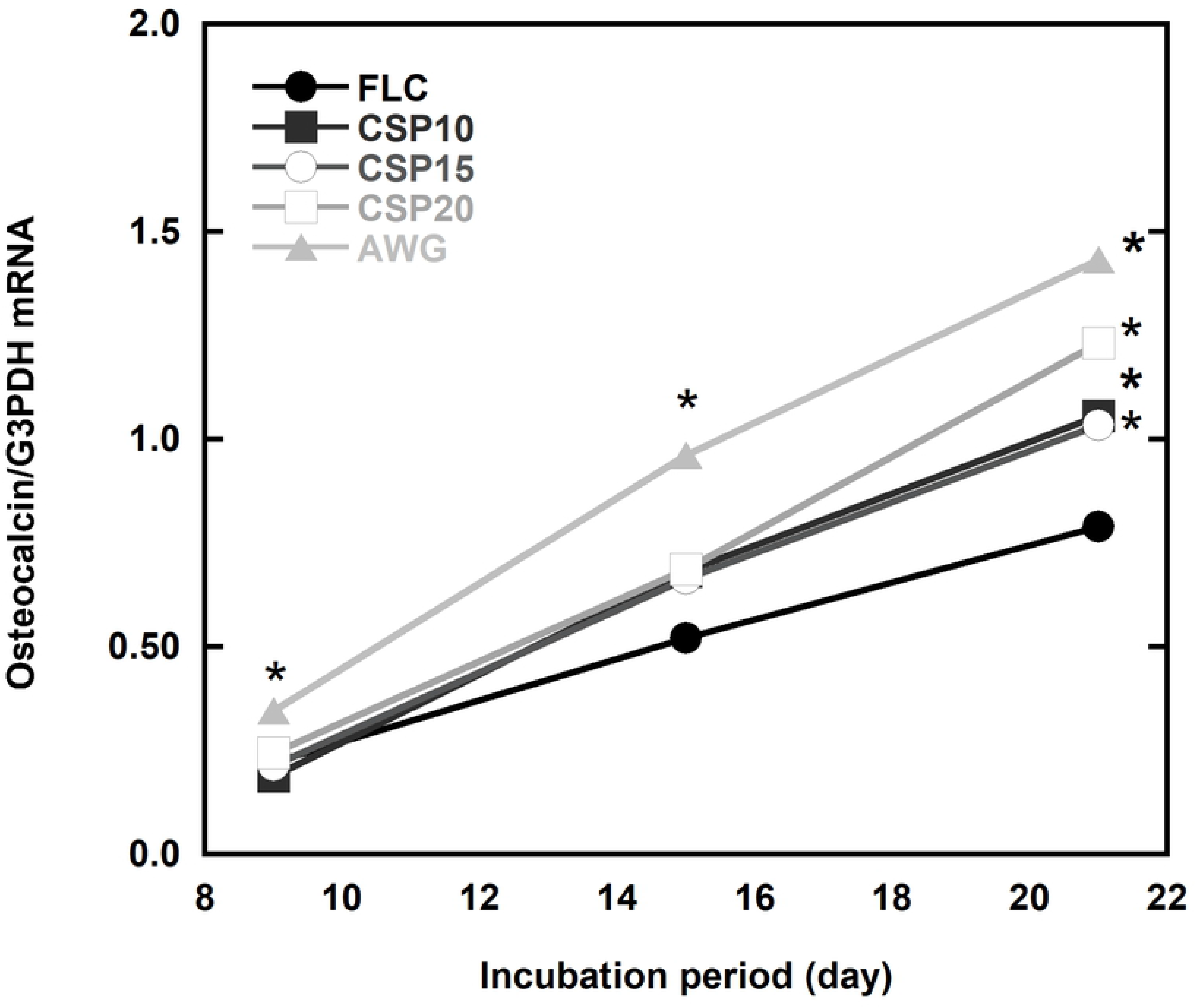
Time course of osteocalcin mRNA expressed by MC3T3E1 on each cement specimens. The results are shown as the mean + standard deviation (n=9). The superscription indicates a significant difference as compared with FLC on the same day.

## Discussion

### Bioactivity on the surface of bioactive cement

Based on surface analysis of the specimens of RMGIC containing BAG, apatite phase was formed after 4 and 8 weeks immersion in SBF. The diffraction patterns were typical for all cements containing BAG investigated in this study in which peaks were developed around 2*θ* of 25.5 and 32.0°. Comparing the patterns with that for a synthetic commercial hydroxyapatite (HAP) as shown in **Figure 2**, the HAP peak around 2*θ* of 32.0° was more prominent, showing higher crystallinity than that of the cement containing BAG. This fact suggested that apatite formed on the surface of RMGIC containing BAG was poorly crystalline [17–18]. This fact also allowed RMGIC, with less contain of polyacrylic acid than conventional GIC, to form an apatite layer on the surface of the cement after immersion in SBF by incorporation of BAG [19].

It was reported that with pure CaO-SiO_2_-P_2_O_5_ glasses and AW glass ceramics, the apatite development was observed after immersion in SBF for only 7 days [12, 20]. The delayed apatite formation in 4-8 weeks immersion observed in the present study was probably due to the ratio of BAG compared to the presence of polyacrylic acid inside the cement. In this study, less amount of BAG incorporated in the RMGIC also found to delay apatite formation.

Detection of the apatite formation on the bioactive RMGIC was confirmed from the spherical apatite layer found from the micrograph of the samples. Representative bioactive ceramics contain hydroxyapatite or its components, such as CaO and P2O5, had been believed to be able to integrate with bone in the body because of apatite-forming ability to act as an interface [21]. However, assessments in SBF related to the apatite forming ability on materials with different compositions imply that CaO and P2O5 are not the essential components for apatite formation [20]. In the CaO-P_2_O_5_-SiO_2_ system it is also known from some studies [21–23] that the composition of the glass forming an apatite layer in an SBF was based on CaO-SiO_2_ system and not CaO-P_2_O_5_ system.

The findings in this study corroborate with the previous studies regarding CaO-SiO_2_ system. It has been clear from the FT-IR spectra that the cements containing CSP10(50) and CSP15(50) have higher ability to form apatite when immersed in SBF with the development of absorption bands around 873, 602 and 568 cm^-1^ assigned to CO3 and PO4 in carbonated apatite besides the asymmetric stretching vibration of the carboxylates at 1563cm^-1^ (**Figure 4**). The increase in intensity of a peak at 1563 cm^-1^ assigned to asymmetric stretching vibration of carboxylate was due to a progress of the acid-base reaction between COOH groups and metal cations like Ca^2+^, Sr^2+^ and Al^3+^ by water absorption in the resin matrix [24]. The glass released Ca^2+^ ion from its surfaces via exchange with the H3O^+^ ion in SBF to form Si-OH groups on their surfaces. The functional Si-OH groups were abundant on the surface of cements containing BAG, especially the ones with CSP10(50) and CSP15(50). This fact explains the preferential formation of the apatite in those cements. The mechanism of carbonated apatite surface layer formation on the resin-modified glass ionomer cements containing bioactive glass could be assumed to be the same as mentioned by Filho and coworkers [25] as described in **Table 4**.

**Table 4.** Infrared frequencies for functional groups on a bioactive glass surface before and after SBF reaction [24].

| Wave number (cm<sup>-1</sup>) | Vibrational Mode |  |
| --- | --- | --- |
| 1350-1080 | P=O | stretch |
| 1250-1100 | P=O | associated |
| 940-860 | Si-O-Si | stretch |
| 890-800 | C-O | stretch |
| 1175-710 | Si-O-Si | tetrahedral |
| 610-600 | P-O | bend crystal |
| 560-550 | P-O | bend amorphous |
| 530-515 | P-O | bend crystal |
| 540-4515 | Si-O-Si | bend |

### Dissolution and precipitation of apatite during the SBF immersion

It was confirmed there was an increase of Ca^2+^ ions concentration in SBF after certain periods of immersions. The Ca^2+^ ions released from the surface of the cement via an exchange with the H_3_O^+^ ion in the SBF and allowed formation of Si-OH on their surfaces [21]. Water molecules in the SBF then or simultaneously react with the Si-O-Si to form additional Si-OH groups. The Si-OH group formed induced apatite nucleation and the released Ca^2+^ ions accelerate apatite nucleation by increasing the ionic activity product (IAP) of apatite in the fluid [20, 22]. Apatite can grow spontaneously by consuming the calcium and phosphate ions in the surrounding fluid once the apatite nuclei are formed, because the simulated body fluid is highly supersaturated with respect to the apatite [20]. This also explained the decrease of phosphate ions by the addition of BAG, since the phosphate released were used to form apatite nuclei.

### F, Sr, and Al ions exchanges and releases

As shown in the results, the fluoride release must have originated entirely from the powder component of the RMGIC. The amount of fluoride releases was also different among the types of the bioactive glasses and the order was different depending on the amount of addition. It is not possible at present to explain such a difference and a further study will be needed. The slopes of the plots were between 0.35 and 0.51 except for CSP10(50), indicating that the fluoride release was determined by the diffusion process through the cement matrix irrespective of the bioactive glass addition. Fluoride release from restorative materials is clinically important because its anti-cariogenicity and the effect in mineralization [26–29]. The result found in this study may indicate that that addition of bioactive glass does not interfere fluoride releasing ability of resin-modified glass ionomer cement.

Meanwhile, Sr release from FLC showed a slope of about 0.44 in the log-log plot and the release rate was controlled by the diffusion mechanism [30–31]. However, the addition of bioactive glasses seemed to alter the mechanism because the slope became larger except for CSP10. The value of slope was about 0.6 and 0.7 with the cements containing 33% and 50% bioactive glasses respectively. This is probably because Sr was involved with the acid-base reaction in the setting process of the cements and Ca from the bioactive glasses altered the role of Sr by participating in the setting reaction.

It was found that the total amount of Al in the cement decreased with the addition of BAG. This fact is closely related to the finding that crosslinking of the carboxylates in the polymeric acid by Al proceeded less in the RMGIC containing BAG as described in the previous study [14, 31]. Addition of the bioactive glasses caused preferential cross-linking by Ca^2+^ ions in the setting process and the Al^3+^ ion remained as unreacted in the original cement powder or free ion in the cement matrix. This may explain the difference in Al release behavior between the cement containing the bioactive glasses and FLC. The slope in the log-log plot was increased to around 1 with the cement containing the BAG. This fact suggests that the Al release was not diffusion controlled but surface reaction controlled [32].

### Bioactivity in the osteoblast like cells (MC3T3E1)

It was confirmed from the cell studies that BAG addition increased bioactivity of the RMGIC significantly. The highest attachment was found on the surface of the CSP10(50) added cement, but there was no difference on the proliferation of the cells among different types of the BAG added. Previous studies demonstrated that surface topology of the substrate could be one among many factors which govern osteoblast attachment. Surface topology represented by surface composition and structure is known to influence the kinetics of protein adsorption and the structure of the adsorbed protein [33–37], thus influence the attachment of osteoblasts. In this study, there were significant differences between FLC and the other set cements that could be explained by the existence of apatite development on the surface of RMGIC containing BAG.

Chang *et al*. [37] reported that the affinity of serum protein components for hydroxyapatite HAP surfaces was so high so that no apparent differences could be found among the three hydroxyapatite surfaces. In this study we found significant difference between cement containing AWG with three other types of RMGIC containing BAG which needs to be investigated further.

Cells differentiation toward bone mineralization was found higher on the cement containing BAG, indicated by type I collagen, ALP, and osteocalcin markers. Ozawa and Kasugai [38] have reported that ALP activity and the number of mineralized nodules on hydroxyapatite produced by rat bone marrow stromal cells were higher than those on a culture plate or titanium. Hot *et al*. [34] have demonstrated that collagen synthesis and calcium uptake by osteoblastic cells were higher on hydroxyapatite than on plastic, and they speculated that the differentiation of osteoblastic cells may be enhanced slightly on hydroxyapatite. Our results obtained in this study with respect to RMGIC containing BAG in comparison with FLC were consistent with their results. In other words, differentiation of osteoblasts was enhanced on the surface of RMGIC containing BAG. The gene expression of type I collagen, ALP, osteocalcin mRNA in the four types of the RMGIC containing BAG were greater than those on FLC.

## Conclusion

It was found that RMGIC containing BAG shows adequate bioactivity by the ability to form carbonate apatite in its surface when immersed in the SBF. The apatite formed was poorly crystalline apatite which has better bone bonding ability. It seems that the addition of BAG to RMGIC did not interfere fluoride releasing ability which is known to be responsible for inhibition of demineralization. Results of Sr and Al analysis lead to the conclusion that addition of bioactive glass may not influence stability of the matrix of RMGIC. It was also confirmed already that Ca release increased by addition of bioactive glass which means positive factor indicating excellent ions exchange with SBF. Furthermore, P concentration which tends to decrease may indicate that apatite formed might grow spontaneously by consuming the Ca and P ions in the surrounding fluid. The ability of osteoblasts differentiation on the four types of resin-modified glass ionomer cement containing bioactive glasses were higher than that of FLC but were the same in all types of RMGIC containing BAG. This may be evidence that all types of the cement containing BAG are osteoconductive. These results may lead to a new insight of the new generation RMGIC-BAG for biomedical purposes.

